# Methylglyoxal mutagenizes single-stranded DNA via Rev1-associated slippage and mispairing

**DOI:** 10.1101/2025.03.18.643935

**Authors:** Sriram Vijayraghavan, Alessandra Ruggiero, Samuel Becker, Piotr Mieczkowski, George S. Hanna, Mark T Hamann, Natalie Saini

**Author notes:** These authors contributed equally to the study (Sriram Vijayraghavan, Alessandra Ruggiero).

## Abstract

Methylglyoxal (MG) is a highly reactive aldehyde that is produced endogenously during metabolism and is derived from exogenous sources such as sugary food items and cigarette smoke. Unless detoxified by glyoxalases (Glo1 and Glo2), MG can readily react with all major biomolecules, including DNA and proteins, generating characteristic lesions and glycation-derived by- products. As a result, MG exposure has been linked to a variety of human diseases, including cancers. Prior studies show that MG can glycate DNA, preferentially on guanine residues, and cause DNA damage. However, the mutagenicity of MG is poorly understood in vivo. In the context of cancer, it is essential to comprehend the true contribution of MG to genome instability and global mutational burden. In the present study, we show that MG can robustly mutagenize induced single-stranded DNA (ssDNA) in yeast, within a guanine centered mutable motif. We demonstrate that genome-wide MG mutagenesis in ssDNA is greatly elevated throughout the genome in the absence of Glo1, and abrogated in the presence of the aldehyde quencher aminoguanidine. We uncovered strand slippage and mispairing as the predominant mechanism for generation of all MG-associated mutations, and demonstrate that the translesion polymerase Rev1 is necessary in this pathway. Finally, we find that the primary MG-associated mutation is enriched in a variety of sequenced tumor datasets. We discuss the genomic impact of methylglyoxal exposure in the context of mutagenesis, DNA damage, and carcinogenesis.

## Main

Methylglyoxal (MG) is a highly reactive dicarbonyl compound. Endogenously, MG is primarily formed from unstable triose phosphate glycolytic intermediates, but MG can also form via lipid peroxidation, ketone body oxidation, and via amino acid catabolism ^1,2^. Further, dietary sources such as sugary foods and flavoring agents, beverages like tea and coffee, tobacco smoke are well-documented exogenous sources of MG (reviewed in ^1^). MG is toxic to cells through its strongly electrophilic nature, which allows it to robustly glycate biomolecules such as proteins, lipids, and DNA ^2–4^. MG-derived advanced glycation end products (AGEs) ^5,6^ are highly detrimental to cellular homeostasis, and as such are linked to numerous human ailments, including neurodegenerative disorders such as Parkinson’s ^7^, diabetes ^8^, cardiovascular diseases and renal diseases ^9,10^, aging ^11^, obesity ^12^, and cancers ^13,14^

In cells, the primary mode of MG detoxification is via the evolutionarily conserved glyoxalase Glo1, which converts MG to S-d-lactoylglutathione via a reduced glutathione cofactor ^15^. As such, Glo1 expression is directly linked to MG toxicity. Yeast cells lacking *GLO1* are highly sensitive to MG ^16,17^. Human cells treated with the Glo1 inhibitor S-*p*-bromobenzylglutathione cyclopentyl diester (BBGC) are sensitive to low concentrations of exogenous MG ^18^. Further, Glo1 copy number amplification has been observed in ∼8% of all human cancers ^19^, indicating high endogenous MG levels in the tumor microenvironments. Elevated MG concentrations can be largely attributed to increased anaerobic glycolysis in tumor cells, termed “Warburg effect” ^20^. Conversely, MG levels can be diminished via MG scavengers such as L-carnosine ^21^ and aminoguanidine ^22^ which prevent MG accumulation, or through the suppression MG formation by targeting glucose metabolism, via drugs such as metformin ^23,24^.

MG primarily reacts with deoxyguanosine residues to form the nucleotide adduct N^2^-(1-carboxyethyl)-deoxyguanosine (CEdG) ^25^ which undergoes mutagenic lesion bypass ^26^. Additionally, MG can induce the formation of covalent DNA interstrand crosslinks (ICLs) ^27^, as well as DNA: protein crosslinks (DPCs) ^28–31^, which are deleterious lesions that can impede normal replication and transcription and induce DNA single and double strand breaks ^32^.

Mutational analyses using *supF* bearing shuttle vectors showed that MG can induce C:G to G:C transversions ^33^. Prevention of MG-associated mutations were primarily seen to be dependent on nucleotide excision repair (NER) in *E.coli* and human fibroblasts ^34–36^. On a genomic scale, MG exposure has been associated with sister chromatid cohesion defects, chromosomal instability, and micronuclei formation in WIL2-NS lymphocyte cell lines ^37,38^.

Even though numerous studies demonstrate the genotoxicity of MG, there is little known about the preferred genomic substrates (single-stranded DNA (ssDNA) or double-stranded DNA (dsDNA)), types of mutations, and mutation signatures associated with methylglyoxal exposure. High fat and sugar western diets are associated with increased adipose tissue dysregulation, diabetic insulin resistance, and represent a significant co-morbidity for metabolic disease, hepatic and gastrointestinal cancers ^39–41^. Physiological changes associated with such diets drastically upregulate MG production in cells. While MG broadly affects multiple molecular pathways, in particular, the genotoxic effects of MG could persist beyond the timeframe of exposure in the form of mutations, and potentially serve as a biomarker for disease onset, progression, and severity. The absence of this information has so far precluded the measurement of MG-contributed mutations in overall disease pathophysiology. Therefore, it is imperative to identify the genetic constraints on MG-associated mutagenesis and suitably characterize the mutational landscape of MG *in vivo*.

Here, we analyze the mutational landscape of MG exposure across the genome. Using budding yeast strains engineered to induce genome-wide single-stranded DNA (ssDNA), we demonstrate that MG is a ssDNA-specific mutagen and results in slippage and mispairing-induced mutagenesis. We further show that the catalytic activity of Rev1 is necessary for MG-induced slippage and mispairing mutagenesis. Finally, we identify MG mutagenesis in sequenced cancer databases, hinting at MG-induced DNA damage across a wide variety of tumors.

## Results

### MG strongly mutagenizes ssDNA in the absence of Glo1 activity

To test MG mutagenicity, we utilized yeast strains harboring a temperature-sensitive *cdc13-1* allele ^42^. When incubated at non-permissive temperature, strains arrest in G2 resulting from global telomere uncapping and 5’→3’ resection, generating large tracts of single stranded DNA ^43,44^. The subsequent addition of mutagens to these strains generates base lesions on ssDNA, which undergo mutagenic bypass upon restoration of permissive growth conditions to allow DNA synthesis to continue. Because of a lack of templated repair of ssDNA lesions, damage is not erased, culminating into somatic mutations. Furthermore, *CAN1* and *ADE2* located on the sub-telomeric left arm of ChrV allow selection of clustered mutations (Figure 1A). Canavanine resistant, Ade-mutants (Can^R^ Ade^−^) appear as red colonies on selective media supplemented with canavanine and low adenine. The selection of double mutations allows to limit background noise for downstream mutation analysis.

**Figure 1:**
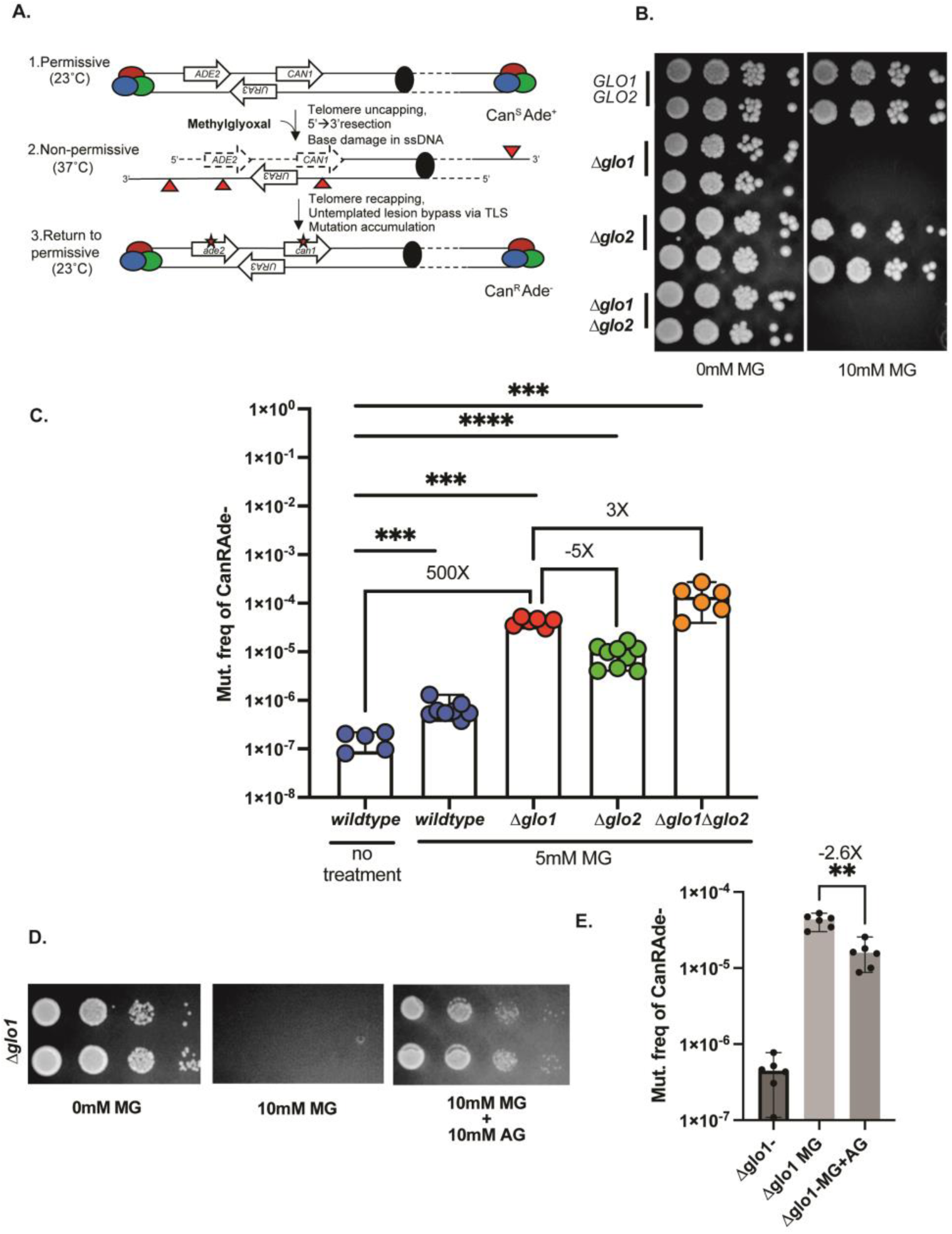
Methylglyoxal is mutagenic on ssDNA. A. Schematic of ssDNA induction and MG exposure. Colored circles represent the telomere capping complex. Solid black circle represents the centromere. Red triangles represent base lesions. Asterisks represent mutations. B. Spot dilutions to test sensitivity of yeast strains to10mM Methylglyoxal (MG). Sloping triangles represent decreasing concentration of cells from left to right. C. Can^R^Ade^−^ mutation frequencies of *glo* deletion strains in response to MG. D. Spot dilutions to test sensitivity of *Δglo1* strains to 10mM MG with or without 10mM aminoguanidine (AG). E. Can^R^Ade^−^ mutation frequencies of *glo1* response to 1hr, 5mM MG treatment with or without 10mM aminoguanidine (AG). Data represents median frequencies with 95% CI. Asterisks represent p-value <0.05 based on a two-tailed unpaired t-test.

Using this system, we observed a 5-fold increase in the frequency of Can^R^ Ade^−^ mutants when wildtype cells were treated with 20mM methylglyoxal. No loss in viability was observed for cultures treated with this concentration of methylglyoxal (Figure S1A, Table S2). In yeast, the primary enzyme that detoxifies MG is the glutathione-dependent glyoxalase Glo1, encoded by the *GLO1* gene ^16,45,46^. Previous studies have shown that ablating *GLO1*greatly increases the sensitivity of cells to exogenous MG ^16,17^. In agreement with prior observations, we noticed that *Δglo1* yeast had markedly reduced viability on plates with much lower concentrations of MG (10mM, Figure 1B, Table S2). We then asked if MG is mutagenic in Glo1-deficient backgrounds. Based on our plate viability assays, we conducted our mutagenesis assays by treating cells with a sub-lethal dose of MG (5mM) for an hour. Compared to untreated wildtype cells, wildtype cells treated with 5mM MG displayed a ∼6-fold increase in Can^R^ Ade^−^ mutation frequencies (Figure 1C, Table S2). In comparison, with Δ*glo1* strains treated with 5mM MG showed a nearly 500-fold increase in mutation frequencies (Figure 1C, Table S2). In cells, Glo2 acts as a backup pathway that predominantly acts to detoxify other oxo-aldehydes such as glyoxal ^47^. Deletion of *GLO2* had no effect on viability in response to exogenous MG (Figure 1B-C, Table S2). *Δglo2* strains exhibited >5-fold lower mutagenesis compared to *Δglo1* strains in the presence of 5mM MG (Figure 1C, Table S2). Finally, *Δglo1Δglo2* double mutants phenocopied *Δglo1* single mutants for viability (Figure 1B), and displayed a >3-fold higher mutagenesis in response to exogenous MG compared to the *Δglo1* mutants treated with 5mM MG (Figure 1 B-C, Table S2). The short duration (1h) of MG treatment did not result in a significant loss of viability for any of the above strains (Figure S1B). Our results demonstrate that MG is highly mutagenic on ssDNA in the absence of functional glyoxylase activity, the latter being principally driven by Glo1.

### Aminoguanidine diminishes MG-associated mutagenesis

Aminoguanidine is a potent scavenger that can react with dicarbonyl molecules through its guanidinium group and readily detoxify them ^48^. The scavenging properties of aminoguanidine have been used to treat prevent MG-associated accumulation of AGEs and the treatment of diseases such as diabetic retinopathy and cardiac fibrosis in mouse models ^49,50^. We leveraged the MG-detoxifying properties of aminoguanidine to ask if it lowers MG-associated mutagenesis. Co-treatment of *Δglo1* strains with 10mM MG and 10mM aminoguanidine restored viability of *Δglo1* strains against MG toxicity (Figure 1D), confirming that aminoguanidine effectively scavenges MG in our yeast system. Importantly, co-treatment of *Δglo1* strains with 5mM MG and 10mM aminoguanidine for 1 hour significantly diminished MG-associated mutagenesis (Figure 1E). Our data indicate that scavengers can effectively prevent the accumulation of MG-derived DNA lesions and thereby readily modulate MG-associated mutagenesis.

### MG predominantly generates ssDNA-associated G-mutations

Next we sought to explore the mutation spectrum of methylglyoxal. To this end, we isolated genomic DNA from 114 clonally expanded *Δglo1* Can^R^Ade^−^ mutants treated with 5mM MG, and performed whole genome sequencing. We further sequenced ∼40 independent Can^R^ mutants from water-treated *Δglo1* isolates as controls, as the mutation frequency in these samples was too low to obtain double (Can^R^Ade^−^) mutants. Finally, we included in our analysis ∼40 independent Can^R^Ade^−^ mutants from 5mM MG+10mM aminoguanidine co-treated *Δglo1* strains (Table S3).

We first asked if single base substitutions increased in response to MG treatment. All mutations present in the original parental strains were removed from the mutant isolates to provide the minimum most accurate base substitution (SBS) calls. Compared to controls, MG-treated samples had a significantly elevated mutation burden (Figure 2A, Table S4, (n=761 from 114 samples)). In contrast, water (n=76 from 39 samples) and MG-aminoguanidine co-treated samples had low mutation loads (n=76 from 38samples, Figure 2A). We then analyzed the genome-wide mutation spectrum of the sequenced samples to determine the most prevalent base changes. As per convention, aggregate base substitutions were calculated by adding the total base changes for a given residue with the corresponding reverse complemented base changes, with the final spectrum represented as pyrimidine (C or T) changes. Overall, mutations were elevated in cytosine residues (C→N/G→N, N=A,T, or G/C) for all MG-treated samples (Figure 2B, Table S4). Water, and aminoguanidine-MG co-treated samples had markedly lower levels of overall C→N (N=A,T, or G) changes. For all samples, the overall mutations in thymine residues (T→N/A→N,N=C, G, or A/T) were low. Among all the observed C→N base changes, MG treatment resulted in the highest frequency of cumulative C→G mutations (i.e sum of C→G and G→C mutations) per isolate per base pair in ssDNA compared to other cytosine mutations (C→G=1.13E-05; C→A= 7.46E-06;C→T=6.74E-06, Table S4)

**Figure 2:**
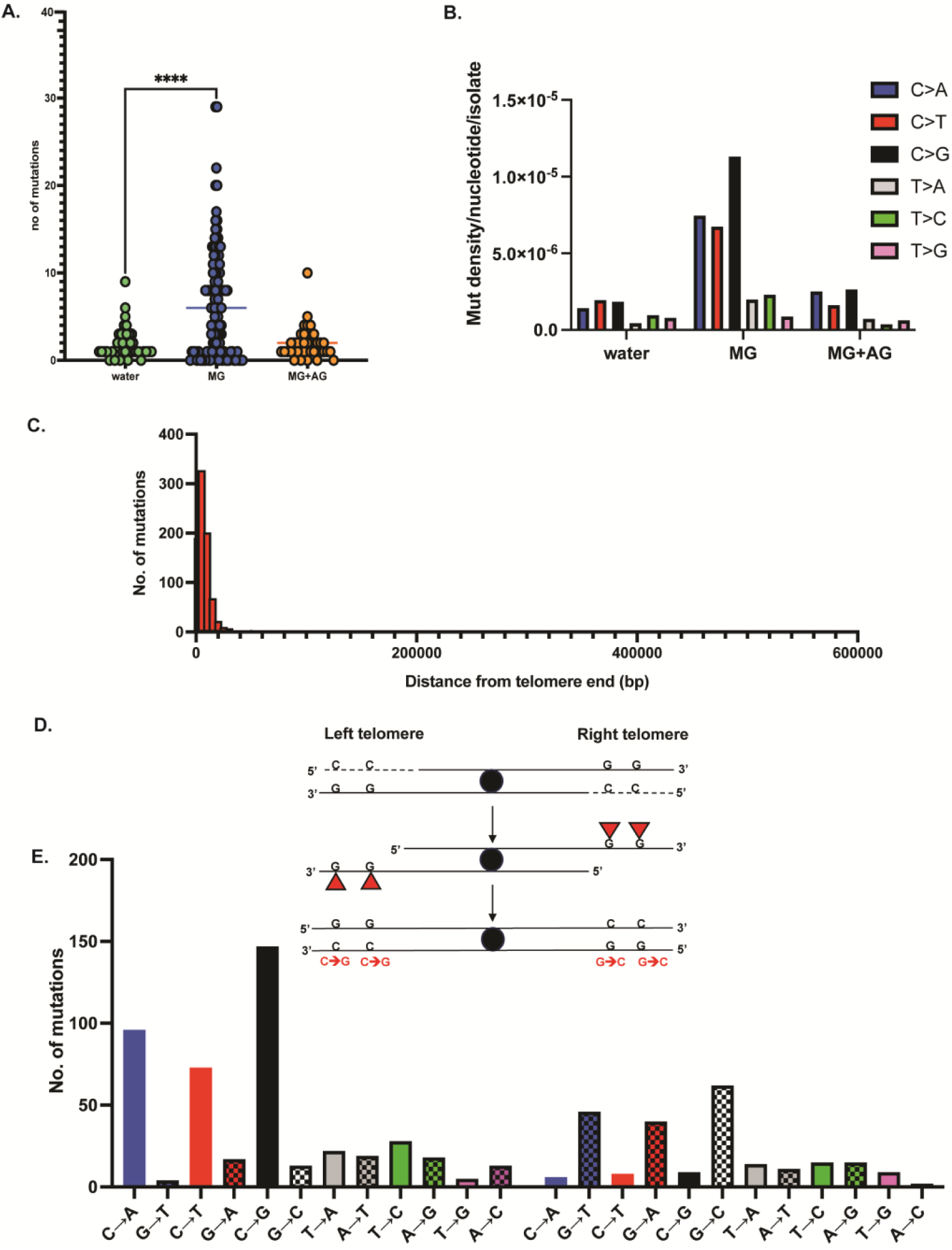
Mutation spectrum of MG on ssDNA. A. Median mutation loads of single base substitutions with no treatment (water), MG, or MG+Aminoguanidine (AG). B. Mutation density of base changes per nucleotide with no treatment (water), MG, or MG+Aminoguanidine (AG). Base changes are cumulative (reverse complement included) but are represented as pyrimidine changes per standard convention. C. Distance of single base substitutions from telomere in MG-treated samples. 0-30000 bp (30kb) represents “sub-telomeric” sequences. Total mutations for all samples are plotted. D. Strand-aware mutational analysis for MG treated samples (top) schematic to display strandedness of mutations, whereby ssDNA lesions are reciprocally reported on different arms of a given chromosome, owing to the direction of resection and sequencing. (bottom) Mutation spectrum of MG treated samples on combined left and right telomeres across all chromosomes. Chromosome coordinates for yeast reference sequence (sacCer3) were obtained from UCSC Table Browser and distances were estimated using BEDtools ^96^.

We subsequently mapped the genomic locations of base changes in the MG-treated samples. In yeast, 5’→3’ resection from telomere ends can extend up to 30 kb, resulting in the accumulation of ssDNA within this region ^51^. We observed that the mutations were predominantly (91% of all base changes) within 30kb from telomere ends across the genome in MG-treated samples (Figure 2C, Table S4). Therefore, MG-associated mutations likely arose from ssDNA-associated lesions. We further sub-classified the sub-telomeric mutations based on their locations on the left vs right sub-telomeric arms. Since 5’→3’ resection would lead to exposed ssDNA on the bottom strand, we expect any base lesions on this strand to be fixed as mutations on the top strand after subsequent replication, thereby making our mutational analysis strand-aware. As such, for any given base change that is enriched on the top strand of the left sub-telomeric regions, the base we expect to observe the reverse complement base substitution to be enriched on the bottom strand of the right sub-telomeric regions (Figure 2D). In line with these predictions, we observed a genome-wide enrichment of C→G changes on the left sub-telomeric regions of all chromosomes and a corresponding increase in G→C changes on the top strand of the right sub-telomeric regions (Figure 2E, Table S4). This strand bias was seen for all mutations in C or G bases, with C>N changes being predominant on the left and G>N changes predominantly on the right. These data indicate that MG predominantly makes lesions/adducts on guanine residues on ssDNA. This is in line with earlier reports that show guanine residues as the major substrate for MG-associated adducts ^3,25,30,33–35,52^. Because the reporter genes involved in mutant selection (*CAN1 ADE2*) were built into the left arm of Chromosome V in our test strains, mutations show a proportional bias towards left telomeres; however, we nevertheless observe a global increase in ssDNA-associated C→N mutations across all telomeres (Table S4). Overall, our SBS analysis demonstrates that the predominant MG-associated mutation spectrum involves cytosine mutations that likely arose from ssDNA-associated guanine base damage.

### MG generates a cCg→G mutation signature

Given the preponderance of cytosine mutations in MG-treated samples, we asked if asked if the mutations displayed any sequence preference around the mutated residue. To investigate this, we used Plogo ^53^ to analyze the context of all major cytosine mutations (C→A,C→T, C→G) in MG- and water-treated samples. Remarkably, for all the above mutation types, we observed that the mutated cytosine copied the base immediately downstream of it (+1 position), thereby generating Cg→Gg, Ct→Tt, and Ca→Aa mutations respectively (mutated residue is capitalized) (Figure 3A). However, based on Plogo, amongst the 3 classes of cytosine mutations, only Cg→G mutations demonstrated a statistically significant over-representation of residue immediately upstream (-1) of the mutated cytosine. We also observed that the mutated cytosine residue was invariably preceded by a cytosine or a faint guanine signal immediately 5’ of it defining the mutation motif as cCg→G (Figure 3A, Table S5). in contrast, no sequence specificity was observed for C→N mutations present in the water-treated samples.

**Figure 3:**
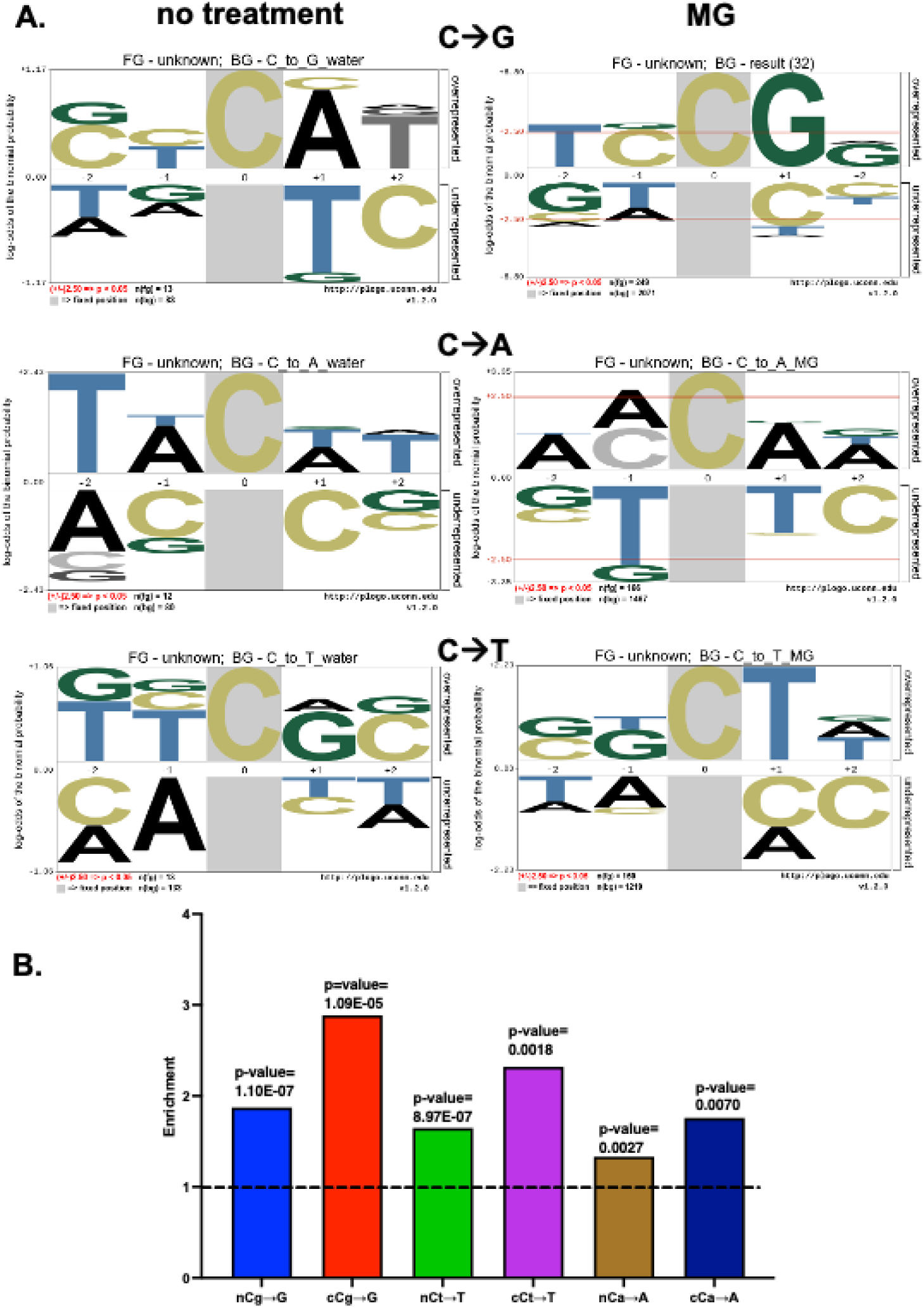
Methylglyoxal mutation signature analysis. A. Plogo analysis of MG mutations. Analysis was carried out for “no-treatment” (water-only controls) samples (left panel) and MG-treated samples (right panel) to measure the over-representation of nucleotides in a pentanucleotide context of C→A, C→T, and C→G mutations. Cytosine in grey highlight represents the fixed C position and heights of residues in the -2 to +2 positions indicate magnitude of over- or under-representation of the indicated residue at the position. N(fg)= foreground mutations i.e total number of C→A substitutions. N(bg)= background mutations i.e number of all other C substitutions across the genome. Red lines in top panel represent over/under-represented residues that are statistically significant. B. Enrichment analysis of MG-treated samples using TriMS ^54^ in various trinucleotide contexts for the predominant base changes C→A, C→T, and C→G. Dotted black line represents the baseline enrichment=1.

To orthogonally determine the mutation motif for methylglyoxal exposure, we used TriMS (Trinucleotide Mutation Signature) analysis ^54^ and asked if any of the above mutational motifs are statistically enriched in MG-treated samples. TriMS predicts the net enrichment and minimum mutation loads for a given base substitution within a trinucleotide context. Specifically, while calculating mutation loads and enrichment, the pipeline corrects for the abundance of the reference base, the mutated base, and the trinucleotide sequence centered on the reference base and the mutated base throughout the genome ^54^. Analysis of mutations in the nC motif versus the cC motif demonstrated that enrichment for cCg→G, cCa→A and cCt→T was greater than nCg→G, nCa→A and nCt→T (Figure 3B, Table S6). cCg→G enrichment was the highest amongst all combinations tested.

These data indicate that methylglyoxal induces adducts on G residues which likely induce blockage of the replicative polymerase, resulting in slippage, mispairing and erroneous copying of the downstream base followed by realignment of the fork and continuation of replication, resulting in cCg→G (cGg→C), cCa→A (tGg→T) and cCt→T (aGg→A) base substitutions (the guanine centered motif is indicated in parentheses).

### MG induces INDELs and multi-base substitutions via slipped strand mispairing

We asked if in addition to single base substitutions, other mutational classes were enriched upon MG treatment. Insertions and deletions (INDELs) were elevated with MG treatment, albeit at a much lower frequency (n=78, median=4 INDELS per isolate, Figure 4) with no increase observed in water-treated samples (n=9, 9 unique isolates). As with SBS mutations, samples co-treated with MG and aminoguanidine had fewer INDELs (n=19, 18 unique isolates), indicating that MG quenching significantly reduces overall mutagenesis (Figure 4A, Table S4). 1-5 bp insertions and deletions were the most-commonly observed tract lengths in MG-treated samples, with insertions mildly outnumbering deletions (Figure 4B, Total insertions=45, total deletions=33, Table S4). Most of the INDELs were present on ChrV (53/78, Table S4), within the left sub-telomeric arm (48/53), likely due to the *CAN1 ADE2* reporter selection bias. Insertions were predominantly at C/G residues (33/45), wherein, A/T were the most inserted bases (28/33). Most insertion events occurred at sites with a G/C residue, indicating that methylglyoxal-induced G-adducts were likely responsible for the insertion events (Table S4). We further investigated the pattern(s) of MG-associated insertions, primarily focusing on the ±20bp sequences around the mutated residue. In striking similarity to the SBS analysis, insertions usually followed the patterns Rm→RMm, whereby R is the reference base, m is the base immediately following the reference base and M is the inserted base identical to m. For instance, almost all the cCa→cAa insertions occurred in CA motifs (Table S4). In contrast, while no specific pattern was observed for MG-associated deletions, ∼40% of deletions occurred in runs of >2 C or G residues (Table S4).

**Figure 4:**
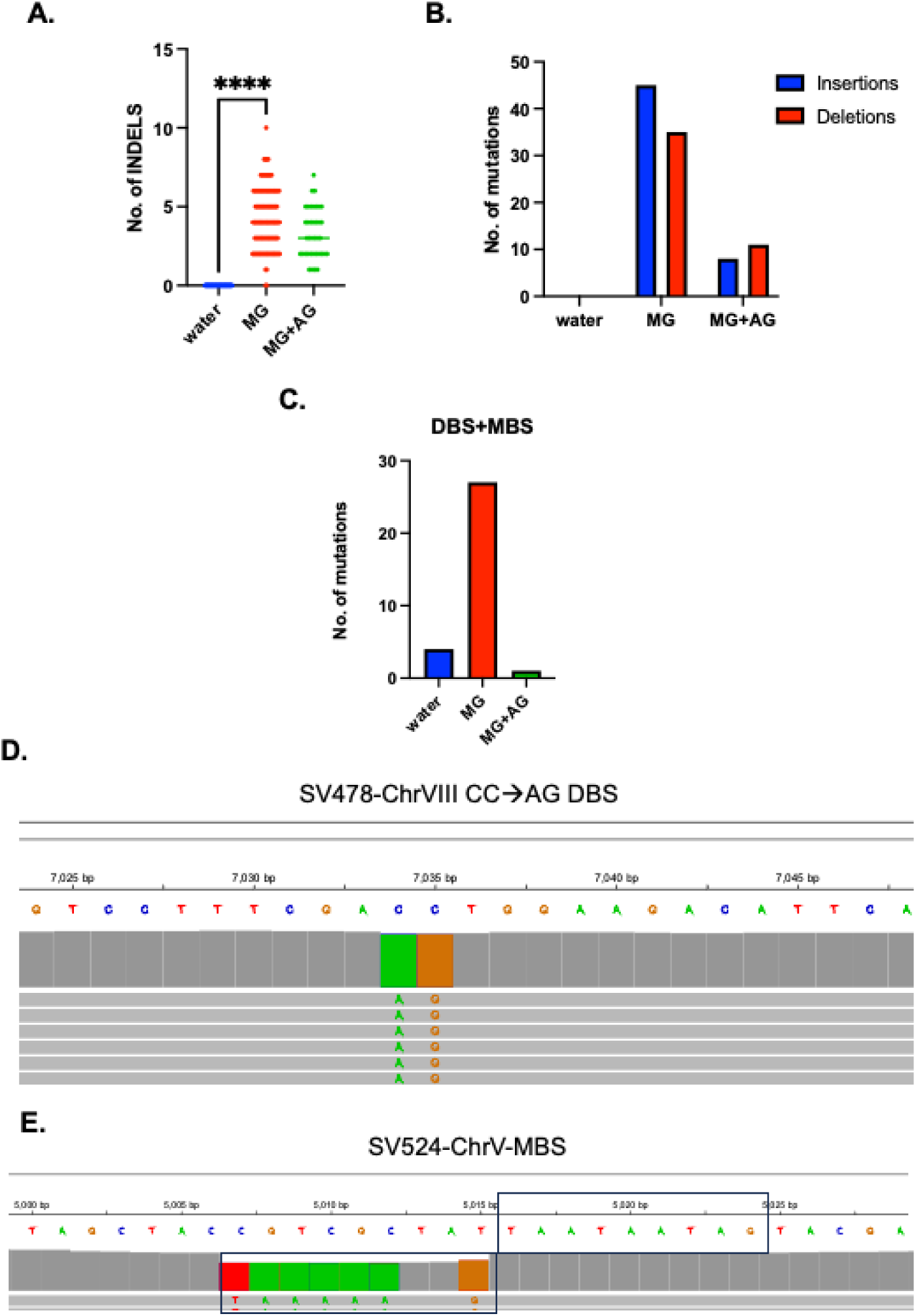
MG exposure induces indels and multi-base substitutions via slippage. A. Median number of INDELS for control (water), MG, and MG+aminoguanidine treated samples. Asterisks represent p-value <0.05 based on an unpaired t-test. B. Proportion of insertions and deletions in the treatment groups from A. C. Cumulative double-(DBS) and multi-base substitutions (MBS) for control (water), MG, and MG+AG treated samples. D and E. Representative examples of MG-associated double-base substitution (panel D) and multi-base substitution (panel E) showing putative template realignment with bases downstream from the reference base and copying. Chromosome plots were generated using the Integrative Genome Viewer (https://igv.org/app/).

In addition, MG-treated samples displayed higher levels of tandem double base (DBS) and multi-base substitutions (MBS) overall compared to control samples (Figure 4C, Table S4). Akin to insertions, we asked if MG-associated DBS and MBS mutations arose via similar slippage, mispairing and copying mechanisms. MG-associated DBS events predominantly occurred on C- or G-containing doublets (19/21,Table S4). Like insertions, tandem base substitutions were templated on consecutive bases within ±10 bp vicinity of the mutated doublets (Figure 4D, Figure S2). Lastly, out of the 5 MBS events, 4 showed tandem base substitutions templated on consecutive bases immediately upstream or downstream of the mutated bases (Figure 4E, Figure S3,Table S4).

Overall, all major MG-associated mutations followed a similar pattern of copying of neighboring bases. This implies that MG-associated lesions on ssDNA-borne guanine residues likely impede replication, and increase the frequency of replication slippage and re-alignment, followed by templated base insertions or substitutions. We conclude that the primary mechanism of MG-associated mutagenesis is slipped strand realignment.

### MG-associated cCg→G mutation bias is eliminated in Rev1-defective strains

In yeast, bypass of ssDNA-associated DNA lesions is generally dependent on three translesion synthesis pathways that utilize low-fidelity DNA polymerases. These comprise of the B-family polymerase Polζ, composed of Rev3 and Rev7 (reviewed in ^55^), the Y-family Pol η polymerase with Rad30 as the catalytic subunit that acts on UV-associated pyrimidine lesions ^56,57^, and the G-template specific polymerase Rev1 that preferentially inserts deoxycytidine residues across abasic sites and coordinates with Rev3 ^58,59^. We asked if either of these factors play a role MG-associated mutagenesis. Removal of *REV3* in a *Δglo1* strain background lowered MG-associated mutagenesis by roughly 18-fold (Figure 5A), demonstrating that MG-associated mutagenesis is heavily dependent on Polζ. Deletion of *RAD30,* which is the catalytic subunit of the error-free TLS polymerase Pol η (Polymerase eta) marginally lowered MG-associated mutagenesis in *Δglo1* strains by <2-fold (Figure 5A). In contrast, we noticed a modest but significant increase (∼2..5X) in MG-associated mutagenesis in *Δglo1* strains harboring a catalytically dead allele of *REV1* (*rev1-aa* ^60^). None of the TLS-deficient strains displayed an appreciable reduction in viability is response to MG-treatment (Figure 5B).

**Figure 5:**
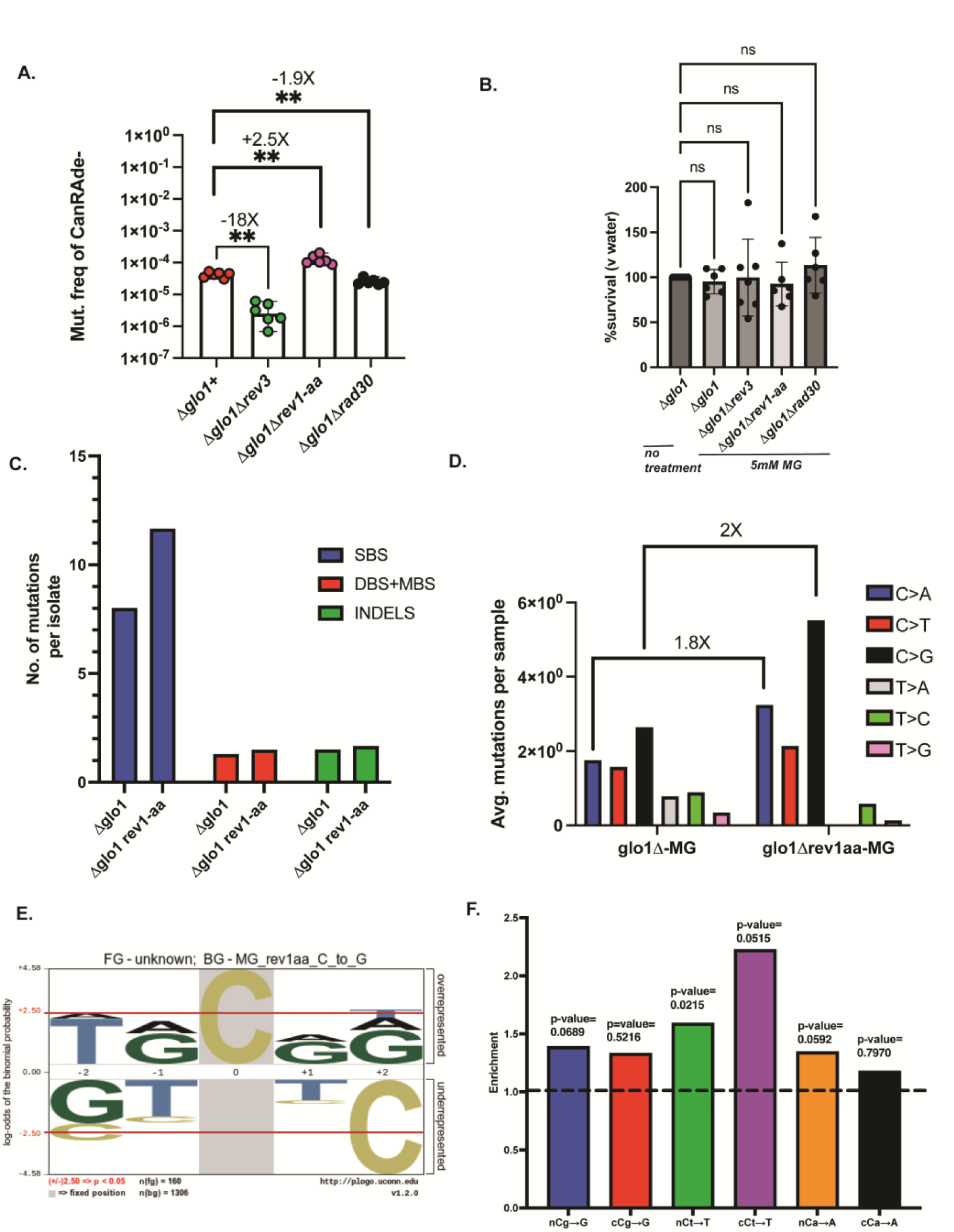
MG-associated mutations are associated with Rev1 activity. A. Can^R^Ade^−^ mutation frequencies of *glo* deletion strains with TLS pathway mutations in response to 1hr, 5mM MG treatment. Data represents median frequencies with 95% CI. Asterisks represent p-value <0.05 based on a two-tailed unpaired t-test B. Viability of strains from panel A in response to 1hr, 5mM MG treatment. Ns-non-significant statistical difference based on an ordinary one-way ANOVA. C. Relative proportions of mutations per isolate in MG-treated *Δglo1* and *Δglo1 rev1-aa* mutant isolates. Only isolates with non-zero mutations are plotted. D. Mutation spectrum of *Δglo1* and *Δglo1 rev1-aa* mutant isolates showing cumulative base changes. Average number of mutations per sample containing non-zero mutations are plotted. E. Plogo analysis of C→G mutations in MG-treated *Δglo1 rev1-aa* isolates. Cytosine in grey highlight represents the fixed C position and heights of residues in the -2 to +2 positions indicate magnitude of over- or under-representation of the indicated residue at the position. N(fg)= foreground mutations i.e total number of C→G substitutions. N(bg)= background mutations i.e number of all other C substitutions across the genome. Red lines in top panel represent over/under-represented residues that are statistically significant. F. Enrichment analysis of MG-treated *Δglo1 rev1-aa* samples using TriMS ^54^ in various trinucleotide contexts for the predominant base changes C→A, C→T, and C→G. Dotted black line represents the baseline enrichment=1.

Because of the observed increase in mutation frequency in the absence of Rev activity, we asked if Rev1-deficient strains had an altered mutation spectrum. To this end, we performed whole-genome sequencing on 30 Can^R^Ade^−^ *Δglo1 rev1-AA* isolates obtained from MG treatment (Table S2), and compared mutations against MG-treated *Δglo1* strains. Overall mutations per isolate were elevated in *Δglo1 rev1-aa* strains compared to *Δglo1* (Figure 5C, Table S7).

Additionally, we saw a marginal increase in the frequency of other mutation types in strains deficient in Rev1 (Figure 5C, Table S7). In comparing the mutation spectra of Rev1 and *rev1-AA* strains, we observed an increase in all cytosine-associated mutations in *Δglo1 rev1-aa* strains (Figure 5D, Table S7). This likely represents the role of Rev1 in accurately bypassing MG-induced lesions by inserting a C opposite the adduct containing G. Interestingly, even though overall C→G mutations were elevated in Rev1-deficient strains, these were no longer enriched within cCg motifs. We confirmed this observation via both Plogo and TriMS (Figure 5 E-F, Table S7). Thus, Rev1 likely has a dual role in MG-associated mutagenesis on guanine residues — directly inserting a correct C opposite some G lesions, while allowing slippage and realignment on a subset of G lesions.

### MG-associated cCg→G mutations are enriched in cancers

To understand the contribution of MG to the overall mutation burden of cancer genomes, we analyzed whole-genome sequenced cancer datasets from the Pan Cancer Atlas of Whole Genomes (PCAWG) ^61^ spanning 1806 samples across 17 cancer types (Table 1), and asked if MG-associated mutations are increased in cancer genomes. Because C→G mutations are the predominant base changes observed upon MG treatment, and cCg appears to be the mutable motif that is combinatorially enriched in both Plogo and TriMS analyses, we infer that cCg→G is the primary mutation motif for MG exposure. Moreover, cCa→A and cCt→T mutations, while enriched with MG treatments, are associated with other etiologies including defective mismatch repair, *POLE* and *POLD* mutations, and tobacco smoke ^62–65^, and can therefore confound further analysis. As such, we focused on cCg→G changes downstream. Using TriMS, we noted that the minimum cCg→G mutation loads were elevated in a wide variety of tumors. Liver (HCC) and lung cancer datasets (LUAD, LUSC) had the highest cCg→G mutation loads, followed by several gynecological tumors (BRCA, OVCA, UCEC) (Figure 6A, Table S8). In addition, an enrichment of the cCg→G signature was observed in at least one sample in several diverse cancers, including biliary tract, stomach, esophageal, head and neck, renal cell, and prostate cancers (Figure 6A, Table S8). We also note that minimum mutation loads for cCg→G changes per tumor ranged from ∼4-89, with HCC, LUAD and LUSC carrying the highest cCg→G mutation loads (Table 1).

**Figure 6:**
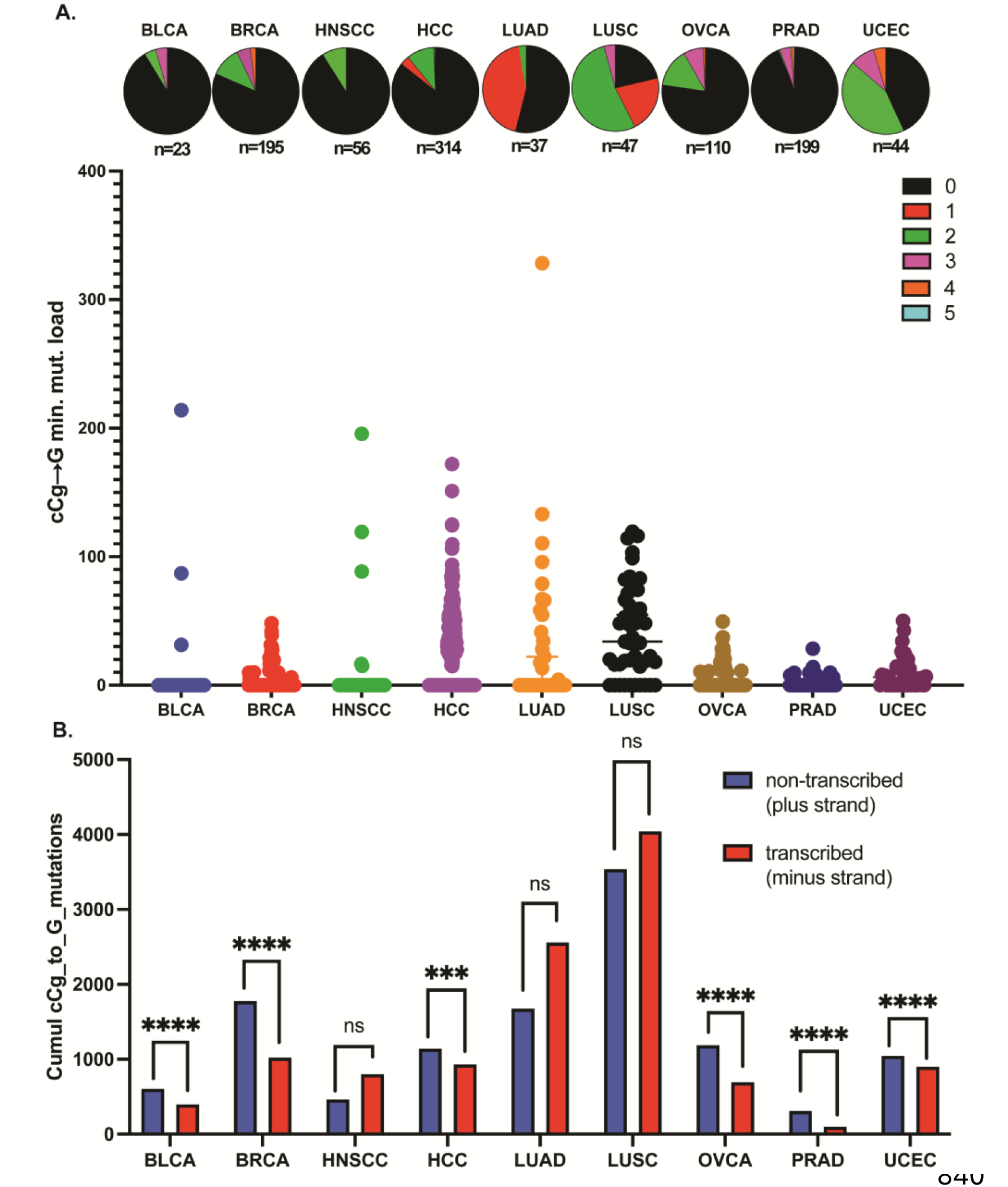
MG-associated mutation signature in whole-genome sequenced PCAWG cancers. A) Scatterplot depicting mutation loads in samples displaying a fold enrichment of the cCg→G mutation signature ≥1 with a Benjamini-Hoechberg corrected p-value of ≤0.05. Samples displaying enrichment are represented as colored sectors in the pie chart situated above the corresponding mutation loads for each cancer dataset. The total number of samples analyzed per cohort is listed under the corresponding pie charts. B. Transcriptional strand bias of the cCg→G mutation signature in PCAWG cancers. Calculations were performed in cancer cohorts displaying a statistically significant fold enrichment of the cCG→G mutation signature in panel A. Benjamini-Hoechberg corrected p-values indicate whether the strand bias is statistically significant. The expanded names for all the abbreviated cancer types are listed. in Table 1

**Table 1:**
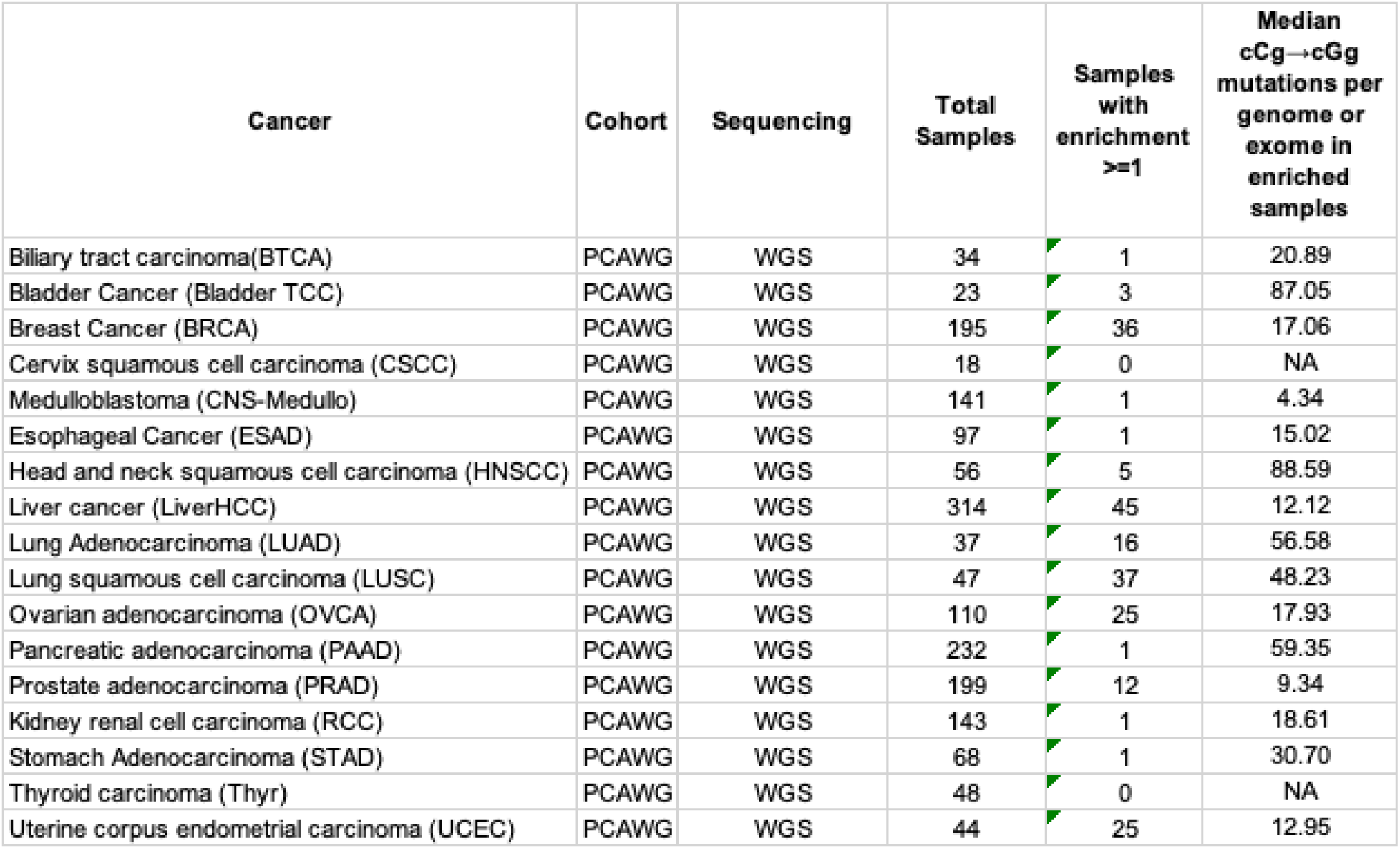
List of cancers analyzed for the cCg→cGg mutation signature. 17 cancer types were analyzed from the PCAWG consortium of whole-genome sequenced cancers. Mean mutation load of the combined cCg→cGg mutations within genomes/exomes were calculated for samples with a statistical enrichment of the cCg→cGg mutation signature (≥1) with a Benjamini-Hoechberg corrected p-value of ≤0.05.

Given that exogenous MG resulted in ssDNA-associated cCg→G mutations in yeast, we hypothesized that ssDNA would be similarly susceptible to MG in cancer cells. To test this, we asked if the cCg→G mutations displayed a transcription-associated strand bias in those cancers where this signature was enriched. The non-transcribed strand in transcription bubbles is single-stranded and therefore susceptible to mutagenesis ^66^. The majority of PCAWG cancer datasets showed a remarkable strand bias for the cCg→G signature towards the non-transcribed strand (Figure 6B, Table S9), strongly suggesting that ssDNA is vulnerable to MG exposure in cancer/pre-cancer cells. However, we presently cannot rule out the role of transcription coupled nucleotide excision repair (TC-NER) in abrogating MG-associated lesions from the transcribed strand, leading to an enrichment of mutations on the non-transcribed strand.

Lastly, given that MG is a mutagenic component of tobacco smoke ^67,68^, we analyzed mutation datasets derived from single cell sequencing of bronchial epithelial cells obtained from subjects with a history of smoking ^69^. In comparison to never-smokers/ex-smokers, cCg→G mutation loads were significantly higher in current smokers (Figure S4C, Table S10). In summary, MG mutation loads were overwhelmingly associated with cancers experiencing elevated MG exposure.

## Discussion

In this study, we demonstrate that methylglyoxal (MG) strongly mutates single-stranded DNA. We show that MG-associated ssDNA mutagenesis is exacerbated in the absence of Glo1-dependent detoxification. In line with prior observations ^2,3,33^, our sequencing analysis reveals that guanine residues are primarily mutated upon exogenous MG exposure, with C→G single base substitutions being the most abundant mutation type. We describe a novel ssDNA-associated mutational motif for methylglyoxal that, to our knowledge, has not yet been ascribed to any other known mutagen, consisting of a cCg/cGg motif. Further, MG-associated mutagenesis relies on Polζ-mediated translesion synthesis, consistent with other studies that demonstrate the role of Polζ in error-prone lesion bypass on untemplated DNA ^55^. MG exposure also gave rise to templated insertions, and multi-base substitutions. Our data suggests that the principal mechanism underlying MG-associated mutagenesis is via strand slippage and realignment, and copying of neighboring bases in a Rev1-dependent fashion. Finally, we observe increased mutation loads of the cCg→G signature across a large and diverse cohort of cancer datasets, suggesting that MG-associated DNA damage and mutagenesis is widespread in tumors.

In our work, we observe an enrichment of C→G (G→C) mutations upon MG treatment. Prior studies using COS-7 and shuttle vector based *supF* reporters have observed a similar mutation spectrum upon treatment with MG ^33^. Such mutagenesis requires the insertion of a G opposite the lesion-carrying G residue. Oxidative damage of guanine bases frequently produces G→T transversions or G→A transitions, but rarely C→G transversions ^70,71^. However, several guanine adducts have the ability to pair with guanine residues at least under *in vitro* conditions ^72^. MG can react with deoxyguanosine in a stoichiometry-dependent manner to generate a variety of adducts with differing stabilities ^52^, and further, MG-guanine adducts can variably pair with A,C, or G residues ^73^. Finally, glycated DNA is chemically different from adducts arising via oxidative damage ^74^. This raises the possibility that *in vivo*, adducts formed upon MG exposure could be subject to differential processing by repair mechanisms. While the short half-life of adducts makes it challenging to predict the base chemistry occurring in the *in vivo* microenvironment, the preponderance of G insertions across a damaged guanine simply suggests that one or more major adducts is a G-pairing lesion, and is enriched on ssDNA upon MG treatment in a concentration-dependent manner. Because MG mutagenicity is largely eliminated in *Δrev3* isolates (Figure 1), adduct templated G insertions are likely translesion synthesis-dependent.

Further, we observed that MG-associated mutations primarily arise via templated realignment and substitutions, whereby bases 3’ of the mutated residue (+1) are copied and substituted at the reference base position. Given that Can^R^Ade^−^ mutation frequencies in *Δglo1* yeast strains were low in *Δrev3* strain backgrounds, we infer that the mechanism of these templated base substitutions and insertions are also TLS dependent bypass of MG-induced mutagenic lesion on guanines. Such slippage and realignment-induced mutagenesis has been previously noted in yeast treated with the platinum chemotherapy agent cisplatin ^75^, and has also been described for the human TLS enzyme Pol Kappa (PolΚ) ^76^. The same mechanism likely extends to longer sequences in the vicinity of the mutated base, which led us to observe templated insertion of multiple bases (2-7 bp) in a subset of the mutations. A similar phenomenon has been previously observed in cisplatin-treated yeast strains, whereby mutations occur via a Rev3-regulated slippage and realignment mechanism ^75^.

We noted that Rev1 catalytic activity was required for the slippage and realignment-induced mutagenesis by MG. When Rev1 is catalytically inactive, we found that C→N substitutions simultaneously go up, but at the same time, the ability to realign template to the neighboring base is also abrogated, leading to disappearance of cCg→G contexts. This might be explained via two co-occurring activities of Rev1. Rev1 is predominantly involved in error free bypass of G-lesions, as it usually puts a C across from an abasic site or an adducted guanine residue ^59,77^.

Presumably, Rev1 initially bypasses MG-adducted guanine residues by incorporating a C opposite the lesion, resulting in error-free damage bypass. However, Rev1 is an inefficient DNA polymerase that can incorporate other nucleotides besides dCTP ^78^. On a subset of G lesions, Rev1 likely copies the downstream nucleotide via base slippage and template realignment, aided by error-prone extension by Rev3 ^79^, leading to C→G, C→A and C→T mutations. Our data also aligns with a prior study that showed an increased sensitivity and mutagenesis of Rev1-deficient yeast strains to another N^2^-dG generating compound 4-NQO, as well as methylglyoxal ^79^, indicating that Rev1 plays a crucial role in survival and mutagenesis in response to MG exposure, both structurally and catalytically.

Overall, our data suggests that repeat sequences, particularly those that are GC-rich, could be susceptible to MG-associated mutagenesis, and subsequent TLS-mediated slippage and realignment-induced errors. It remains possible that MG could induce the formation of intra-strand crosslinks (ICLs) between neighboring GG bases on ssDNA, which are toxic lesions that would likely facilitate polymerase slippage and mispairing. The nature of the adduct and the role of proteins involved in various repair pathways in modulating MG-associated DNA damage and mutagenesis would be worth exploring in future studies.

Most cancers are metabolically dysregulated, which impacts cellular homeostasis via multiple mechanisms. Of note, cells in the hypoxic tumor microenvironments predominantly rely upon glycolytic sugar metabolism for energy production, which is a principal source of methylglyoxal. High MG levels can be cytotoxic, with several studies reporting increased MG-induced cellular apoptosis and inhibition of cancer cell growth *in vitro* ^80–82^. Conversely, sub-toxic levels of MG can promote carcinogenesis, for example, in promotion of metastasis of breast tumors ^83^.

Further, MG adducts have been identified in various cancers, including lung, liver, breast, and skin cancers ^80,84–86^. Additionally, lipid peroxidation is altered in cancer, potentially impacting intracellular MG levels ^87^. Overall, these data demonstrate that MG is prevalent in multiple types of cancers and can be associated with cancer development and progression. In agreement with prior studies, the highest proportion of MG-associated mutations were observed in liver, lung, and breast cancer datasets. We posit that the tissue type, local Glo1 expression levels, and different metabolic and signaling pathways interact in a complex manner to regulate the levels of methylglyoxal in different cancer types.

In lung and esophageal cancer datasets, samples from smokers display significantly higher mutation loads compared to non-smokers (Figure S4 A-B, Table S10). Studies suggest variable concentrations of MG associated with cigarette smoke; 6-60µg MG is present in a single tobacco-based cigarette ^88^, and ∼ 4000-15,000µg/m^3^ MG is present in e-cigarettes ^89^. As such, genomes within cells of the bronchial system likely experience extremely high MG exposure in smokers. In support of this assumption, We believe the observed mutation loads reflect chronic exposure of genomes in lung samples to elevated concentrations of exogenous methylglyoxal.

We also observed a transcriptional strand bias for MG-associated mutations in a majority of the PCAWG cancer datasets, whereby the cumulative cCg→G mutation loads were higher on the non-transcribed/coding strand (Figure 6, Table S9). Based on our yeast data, these data likely reflect a strong propensity for MG to damage ssDNA and/or the inefficient repair of lesions on the non-transcribed strand by the transcription-coupled nucleotide excision repair (TC-NER) ^90^. Interestingly, we did not note a transcriptional strand bias for MG mutations in lung cancers. This might be due to other tobacco-smoke components that have overlapping mutagenic motifs, confounding our analysis.

### Concluding remarks

MG has multiple targets within cells, including proteins, lipids and DNA. The MG mutational signature could either be a result of a direct MG-associated lesion on DNA bases or an indirect consequence of altered protein homeostasis, increased ROS production and subsequent DNA damage, or other types of DNA:protein linkages. At present, we cannot distinguish between these possibilities. In cancer cells, an MG-associated signature may well represent a combination of some or all the above pathways. Given the ubiquity of MG across various tissue types and its association with multiple human ailments, including cancer, identifying and characterizing novel molecular signatures of MG exposure enables a more precise understanding of its role in disease origin and evolution.

## Methods

### Yeast strains

Strains were derived from CG379 with the genotype *MATα his7-2 leu2-3,112 trp1-289, cdc13-1*. The triple reporter strain was constructed as described earlier ^91^. Briefly, *CAN1, URA3, ADE2 and LYS2* were deleted from their original loci and reintroduced as the triple reporter tandem array *lys2::ADE2-URA3-CAN1* on the left arm of Chromosome V at the *de novo* telomere. All gene deletions were made using standard one-step PCR-based methods with dominant drug resistance cassettes *KANMX* or *HPHMX*. The strain carrying the *rev1-AA* allele was the same as described earlier ^92^. All yeast strains and PCR oligonucleotides used in the study are listed in Table S1.

### Methylglyoxal sensitivity and mutagenesis assays

All spot dilution assays were conducted with strains in Table S1 using a 10-fold serial dilution series followed by plating on YPD with or without the indicated concentrations of methylglyoxal ((MG), Millipore Sigma) and/or Aminoguanidine (Millipore Sigma). Spots were incubated for 2-4 days and imaged. For mutagenesis assays, assays were conducted as described previously ^54^, with modifications. Briefly, cultures of the *cdc13-1* strains were grown at 23 °C for 72 hours. Roughly 10^7^ cells were inoculated into fresh YPD and grown with shaking at 37 °C for 4-6 hours in Erlenmeyer flasks to induce G2 arrest from resection at telomeres. Cultures were monitored for complete G2 arrest by analyzing budding index (>95% of cells arrested as large double buds). Thereafter, cells were harvested by centrifugation, washed three times with sterile water and resuspended in water in 15ml conical tubes. MG was added to samples at a final concentration of 5mM and samples were incubated alongside the control samples (without MG) at 37 °C in a rotary shaker for 1h. Dilutions were plated on complete synthetic complete (SC) media (MP Biomedicals) to measure viability and SC-Arginine plates containing 60mg/ml canavanine (Millipore Sigma) and 20mg/ml adenine to isolate Can^R^Ade^−^ mutants (red colonies). All assays with aminoguanidine included 10mM aminoguanidine hydrochloride (Millipore Sigma) in addition to 5mM MG. All plates were incubated at 23 °C for 5-7 days until countable colonies were observed. Median Can^R^ and/or Can^R^Ade^−^ mutation frequencies were calculated as described previously ^54^. Colony counts and image acquisitions were performed using the aCOLyte 3 Automated Colony Counter (Synbiosis Inc.).

### DNA sequencing

Genomic DNA was isolated from independent yeast strains using the Zymo YeastStar genomic DNA isolation kit (Genesee Scientific) per the manufacturer’s Protocol I. DNA was quantified via Qubit (Invitrogen) and diluted to approximately 10ng/µl for library preparation via the Watchmaker DNA library preparation kit (Watchmaker Genomic Inc.) with fragmentation, with each sample acquiring a unique dual index adapter. Illumina NovaSeq6000 sequencing system was used for analysis of pooled libraries.

### Mutation spectrum and signature analysis

Mutation analysis was done as previously described ^54^. Raw sequencing reads were aligned to the reference genome ySR127 ^93^ using BWA-mem ^94^ and duplicate reads were removed using Picard tools (http://broadinstitute.github.io/picard/). Single nucleotide variants (SNVs), insertions-deletions (InDels), double-,and multi-base substitutions were identified using VarScan2 ^95^, using a variant allele frequency filter of 90%. Unique SNVs were by identified by comparing MG-treated samples with untreated parent strains serving as matched normal and after removing duplicates. Mutations were classified as “sub-telomeric” or “mid-chromosomal” based on computed genomic distances from the nearest telomere end using *bedtools closest* ^96^. The cumulative mutation spectra was plotted as pyrimidine changes, taking into consideration reverse complements for every substitution. Mutation strandedness was calculated based on whether the SNVs were located on ssDNA generated upon telomere uncapping and resection. Mutations per isolate were calculated by plotting SNV as a function of the total number of strains used per treatment condition. PLogo ^53^ was used as described previously ^54^ to evaluate the statistical probability of over-/under-representation of residues in the ±1 trinucleotide context of the mutated residue compared to the background sequence. For double-, and multi-base substitutions and indels, nearby genomic contexts were visualized using the Integrative Genomics Viewer ^97^.

### Mutation enrichment and mutation load analysis

Mutation enrichment and mutation loads were calculated based on ^91,92^ using Trinucleotide Mutation Signatures (TriMS) as described previously ^54^. Briefly, total instances of a given substitution in a specific trinucleotide context is compared against its genomewide frequency, as well the incidence of the mutated residue within the ±20 nucleotide context of the mutation. The following calculation was used:

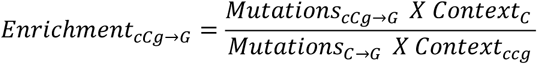

A one-sided Fisher’s Exact test was used to calculate the p-values of enrichment of the given mutation signature in each sample and in the total yeast samples. Mutation loads for a given signature were calculated with a minimum enrichment probability of >1 and a Bonferroni corrected p-value of ≥0.05, using the following equation:

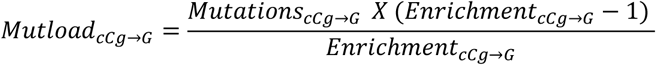

### Mutational analysis in cancers

Somatic mutation load and enrichment were calculated for a given signature using mutation data from de-duplicated somatic SNV calls from different donors in whole-genome-sequenced cancers from PCAWG ^61^ and whole-exome-sequenced cancers from ICGC data portal ^98^. For PCAWG cancer samples carrying an enrichment of the MG-associated mutation signature of ≥1 (Bonferroni-corrected p-value of ≤0.05), transcriptional strand bias of mutations was calculated with BEDTools ^96^ intersect, using hg19 as the reference genome (UCSC Table Browser ^99^) and a goodness of fit test was performed to test the statistical significance of the ratios of mutations on transcribed vs non-transcribed strands using RScript. Smoking metadata from lung cancers was derived from PCAWG ^61^. For analysis of mutation loads in non-cancer bronchial epithelium were obtained from Yoshida *et.al.* ^69^

### Statistics

All statistical tests were performed using Prism V10 (GraphPad Inc.) and RScript.

## Data and code availability

Raw FASTQ sequence files from whole-genome sequencing of yeast samples have been deposited to the Sequence Read Archives (SRA) database under BioProject ID PRJNA1195887. Sequence for the reference yeast genome used in this study (ySR127) is accessible on GenBank (CP011547-CP011563). Source code for TriMS is publicly available on GitHub at https://github.com/SainiLabMUSC/TriMS and is deposited with Zenodo (DOI: https://doi.org/10.5281/zenodo.13862689). The yeast strains used in the study are available upon request.

## Acknowledgements

We would like to thank members of the Saini lab for their critical reading of the manuscript.

## Author contributions

S.V, N.S, M.T.H, G.S.H conceptualized the study. S.V, N.S designed the experiments, S.V, A.R, S.B performed the experiments, P.M performed whole-genome sequencing of yeast samples, A.R., S.V acquired the data, S.V, N.S analyzed the data. S.V and N.S wrote the manuscript. All authors reviewed the manuscript.

---

**Figure S1.**
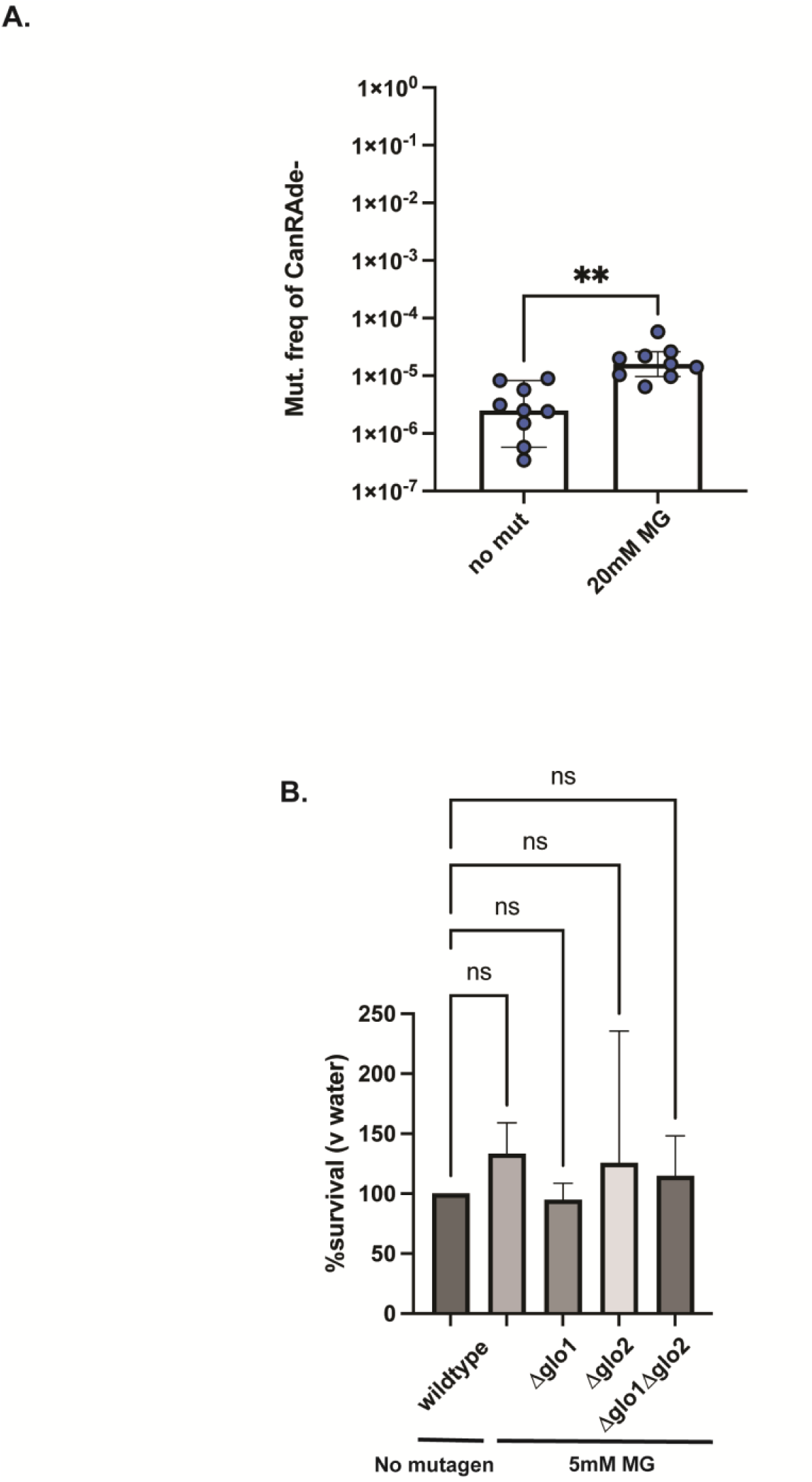
A.Can^R^Ade^−^ mutation frequencies of wildtype strains in response to 1hr, 20mM MG treatment. ** represents a statistically significant difference in median frequencies, indicating a p-value <=0.005 based on an unpaired two-tailed Student’s t-test. B. Viability of strains with MG treatment. Ns-non-significant based on an ordinary one-way ANOVA

**Figure S2.**
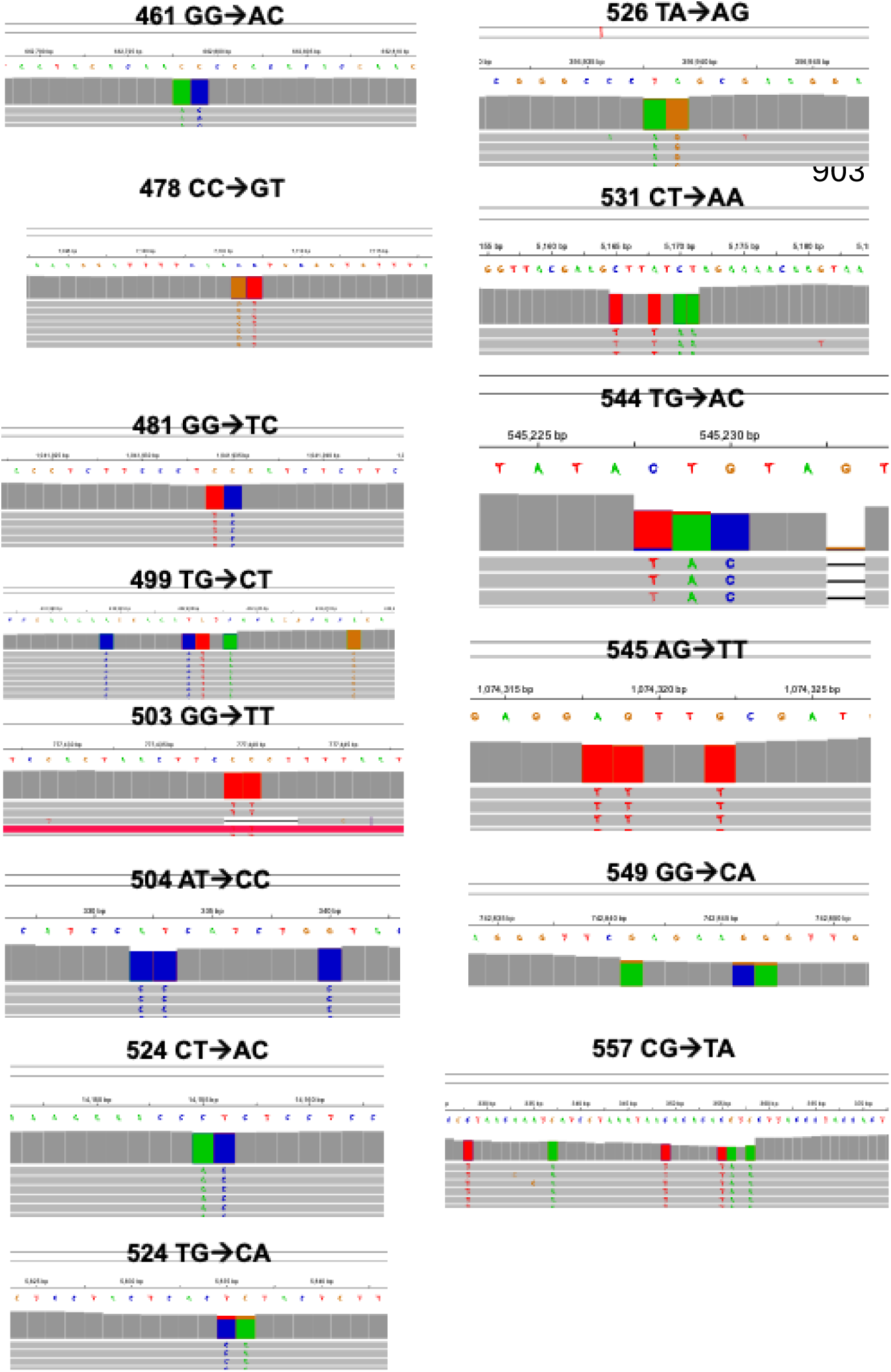
MG-associated double-base substitutions showing putative template realignment with bases downstream from the reference base and the resulting DBS denoted over each plot. Chromosome plots were generated using the Integrative Genome Viewer (https://igv.org/app/).

**Figure S3.**
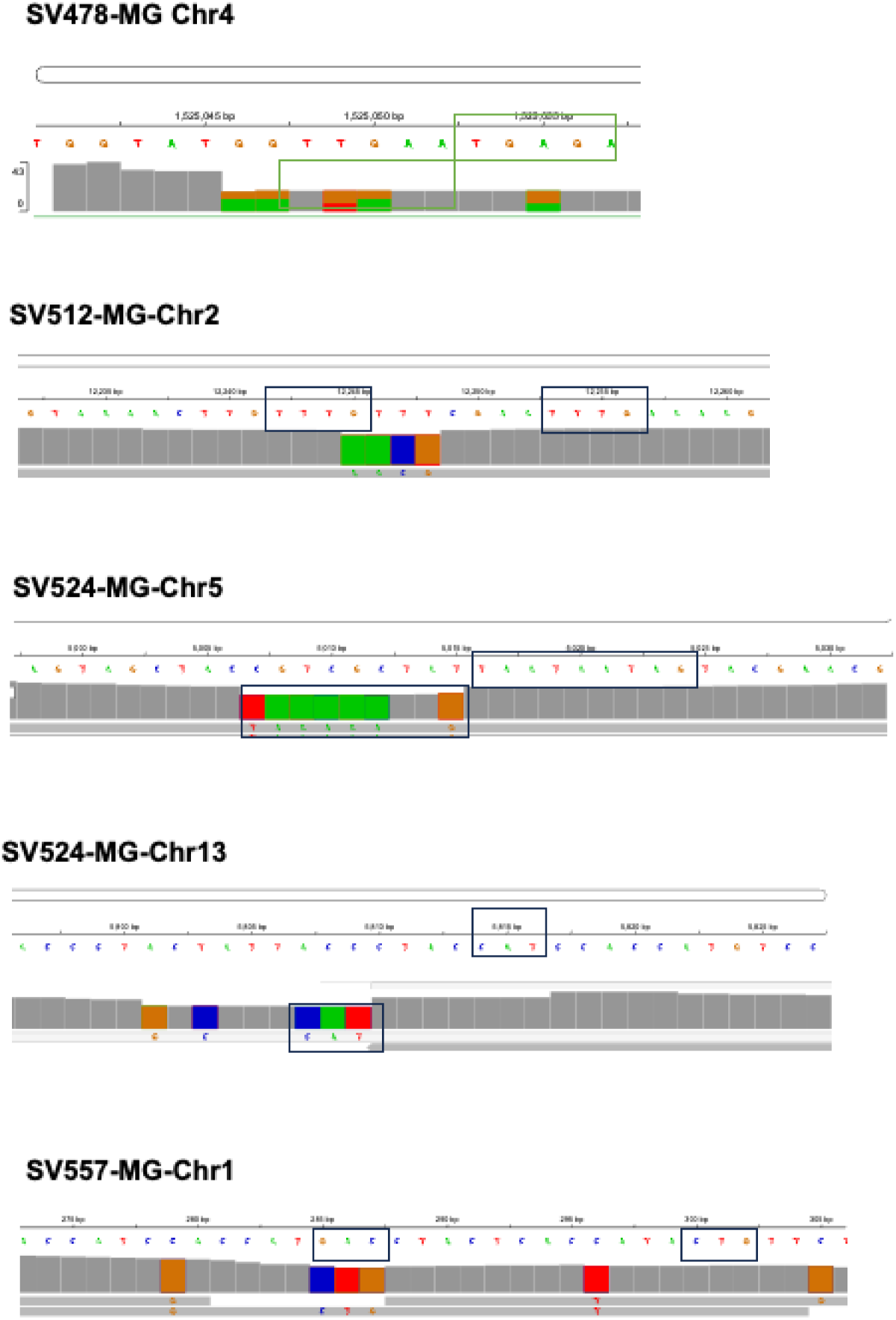
MG-associated multi-base substitutions showing putative template realignment with bases downstream from the reference base and copying. Chromosome plots were generated using the Integrative Genome Viewer (https://igv.org/app/).

**Figure S4.**
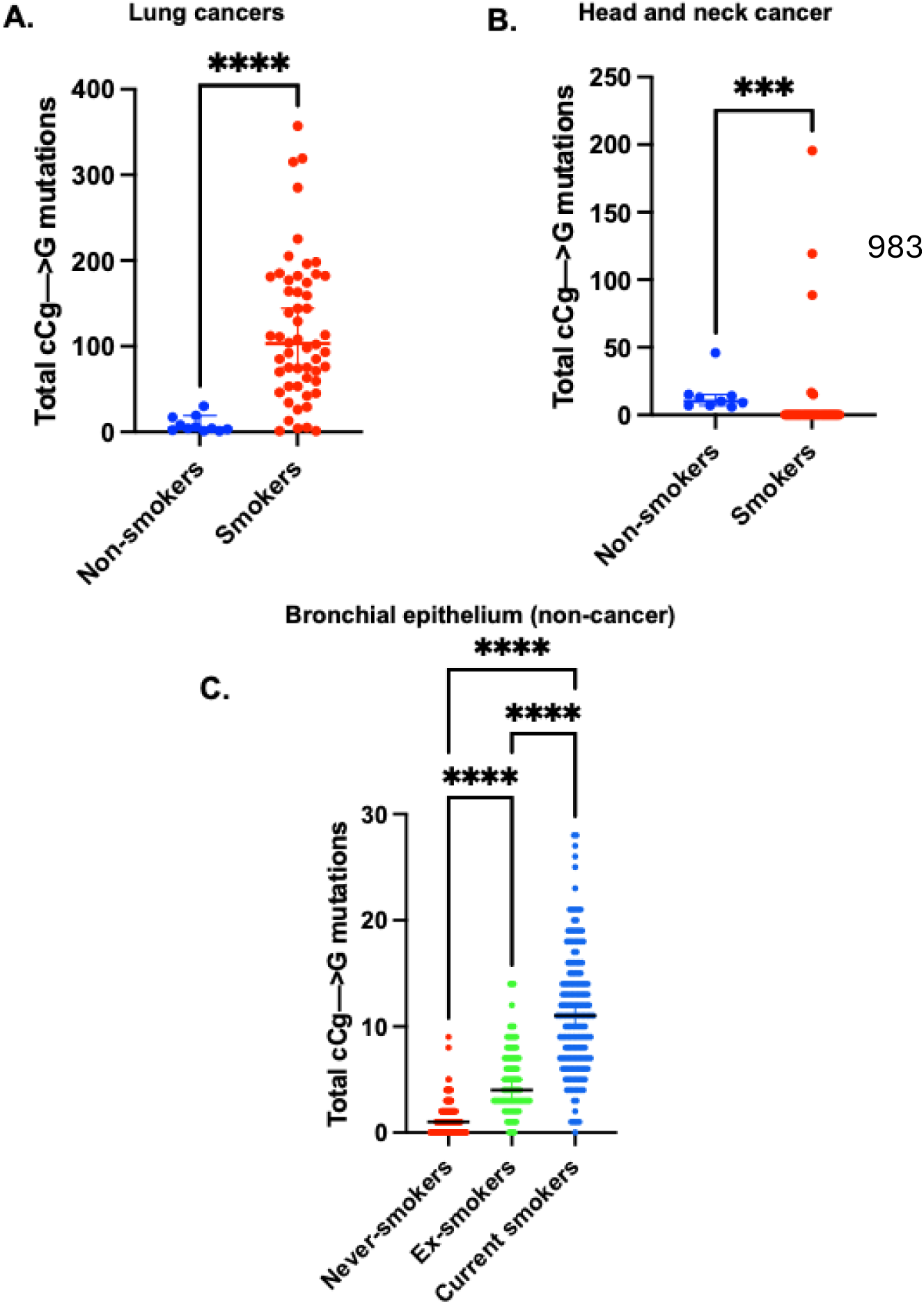
Correlation of cCg→G signature with smoking. A. cCg→G mutations in combined LUAD and LUSC datasets from PCAWG stratified according to smoking status. B. cCg→G mutations in HNSCC data from PCAWG stratified according to smoking status. For both A and B, smoking metadata was obtained from PCAWG. C. Analysis of cCg→G mutation loads in single-cell sequenced datasets from bronchial epithelia of current-, ex-, and never-smokers. Mutation calls and metadata was obtained from ^69^. Asterisks represent p-value <0.05 based on an unpaired t-test.

