## Supplementary figures and images for "Methylglyoxal mutagenizes single-stranded DNA via Rev1-associated slippage and mispairing"

### Figure S1

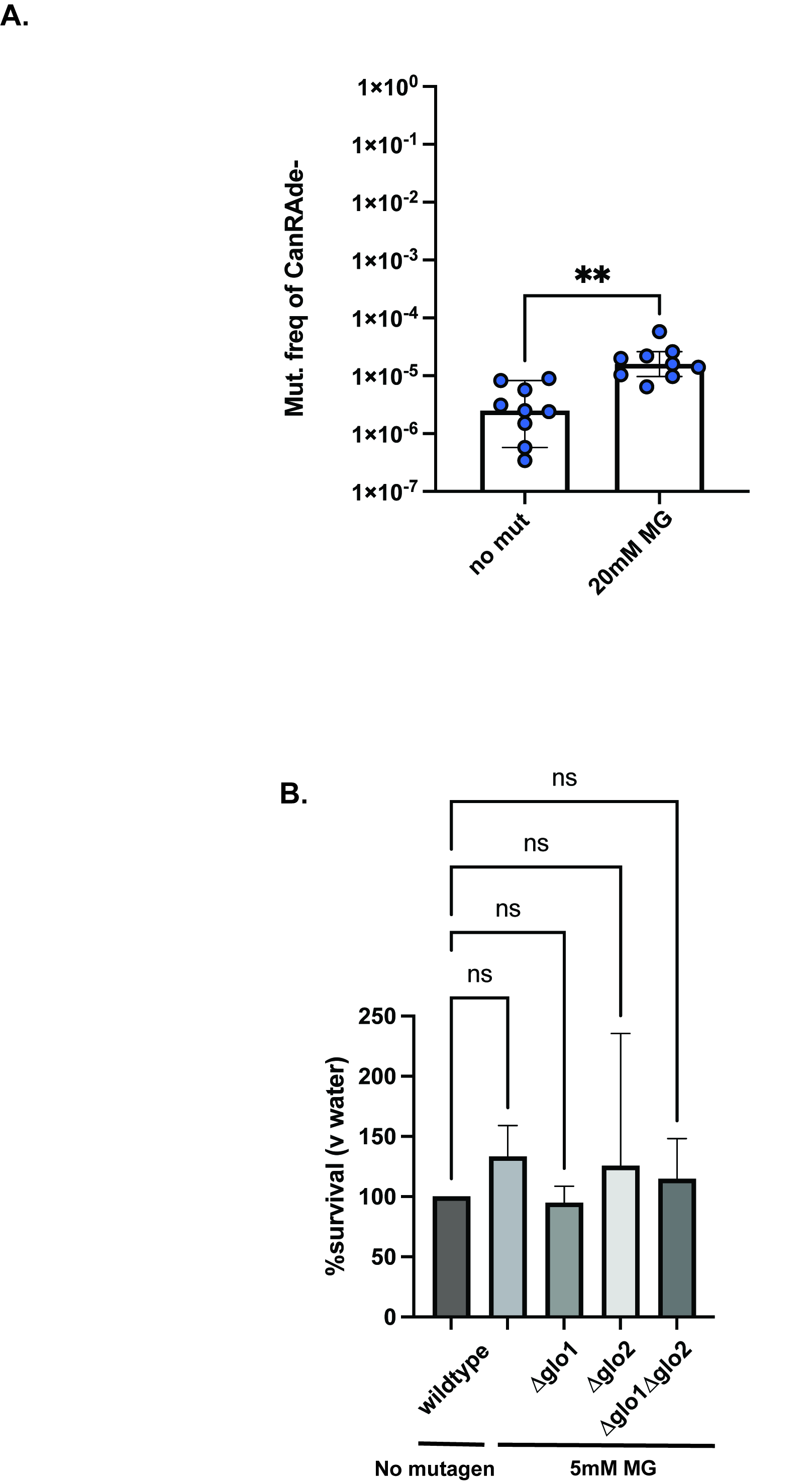

### Figure S2

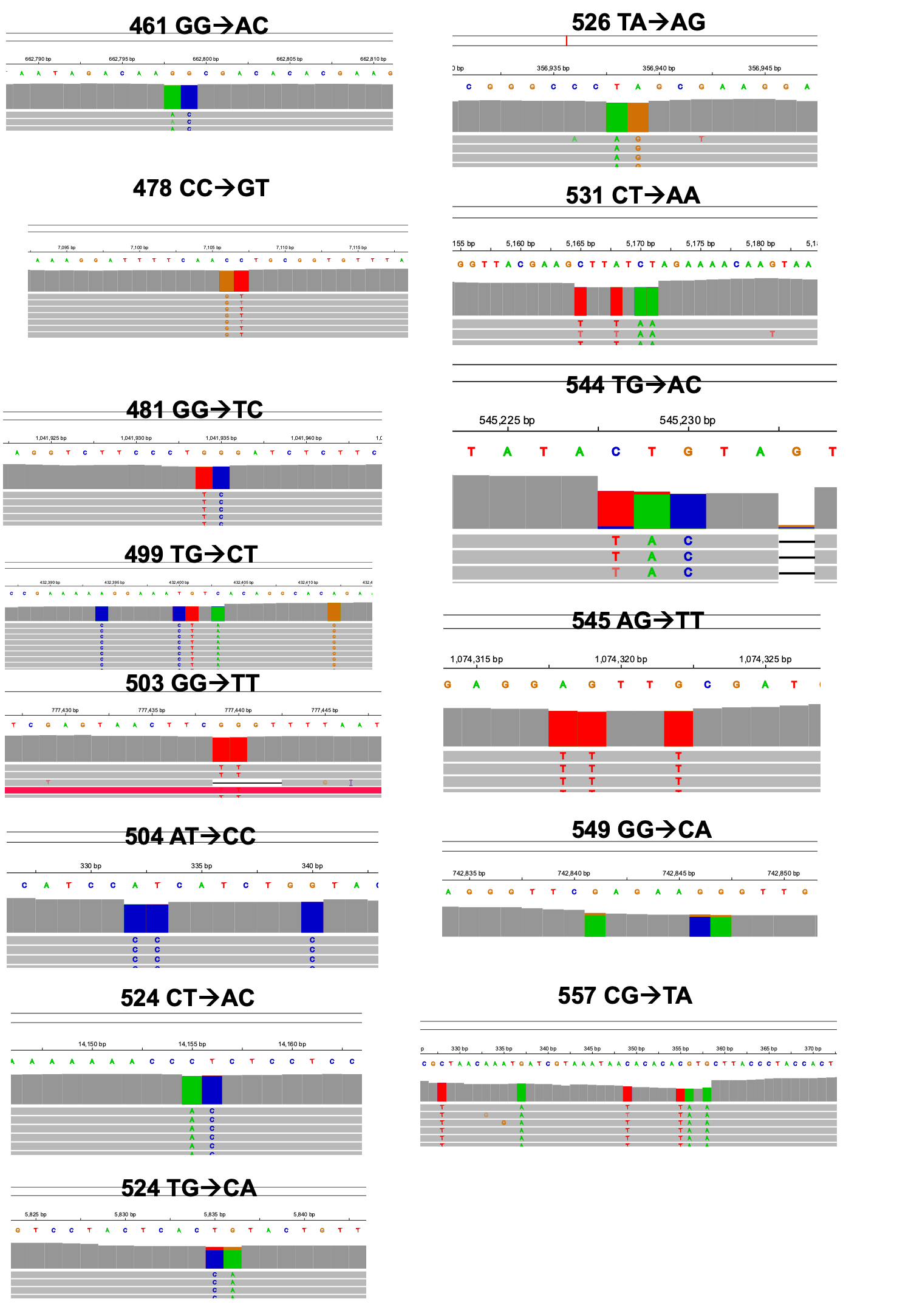

### Figure S3

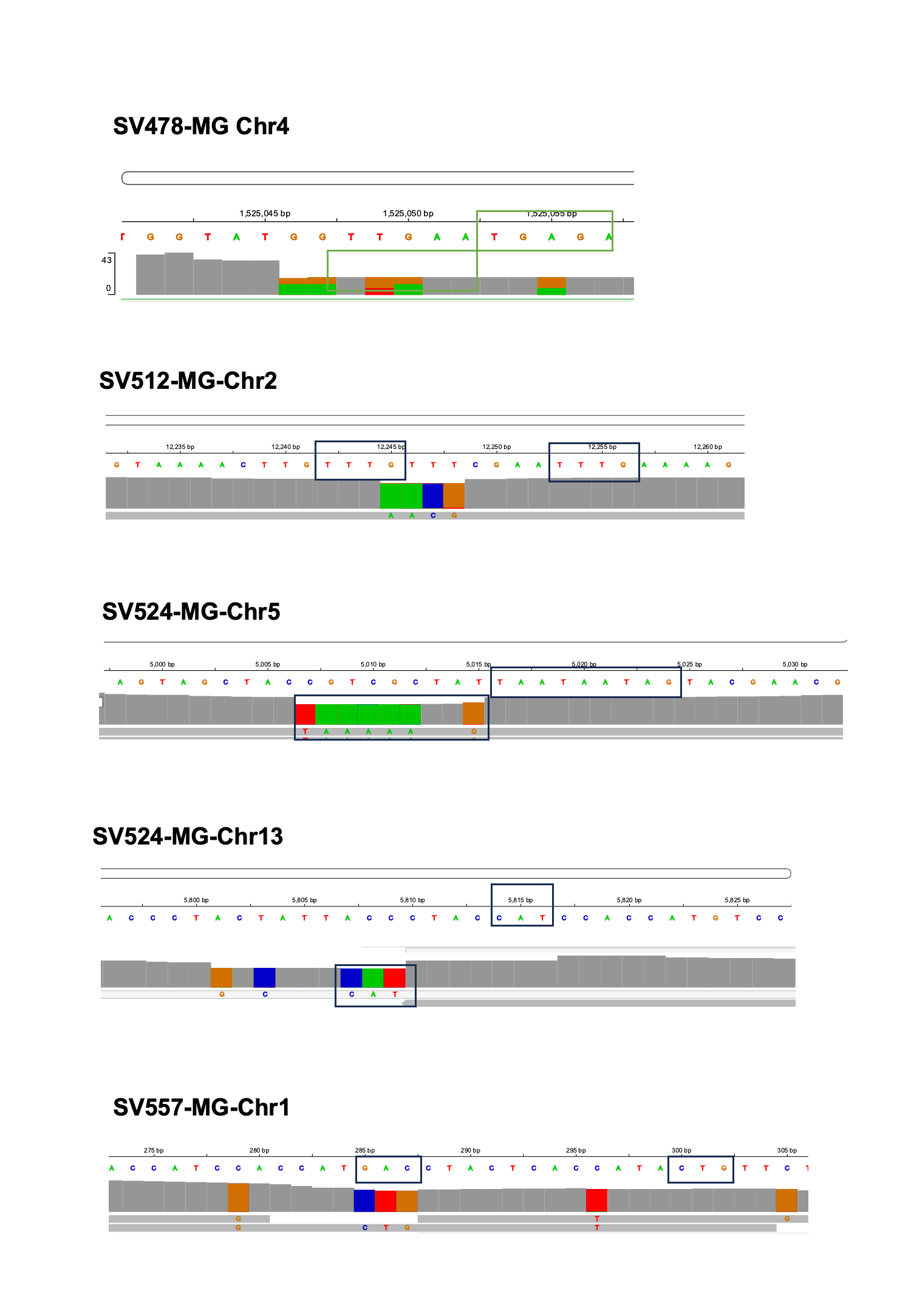

### Figure S4

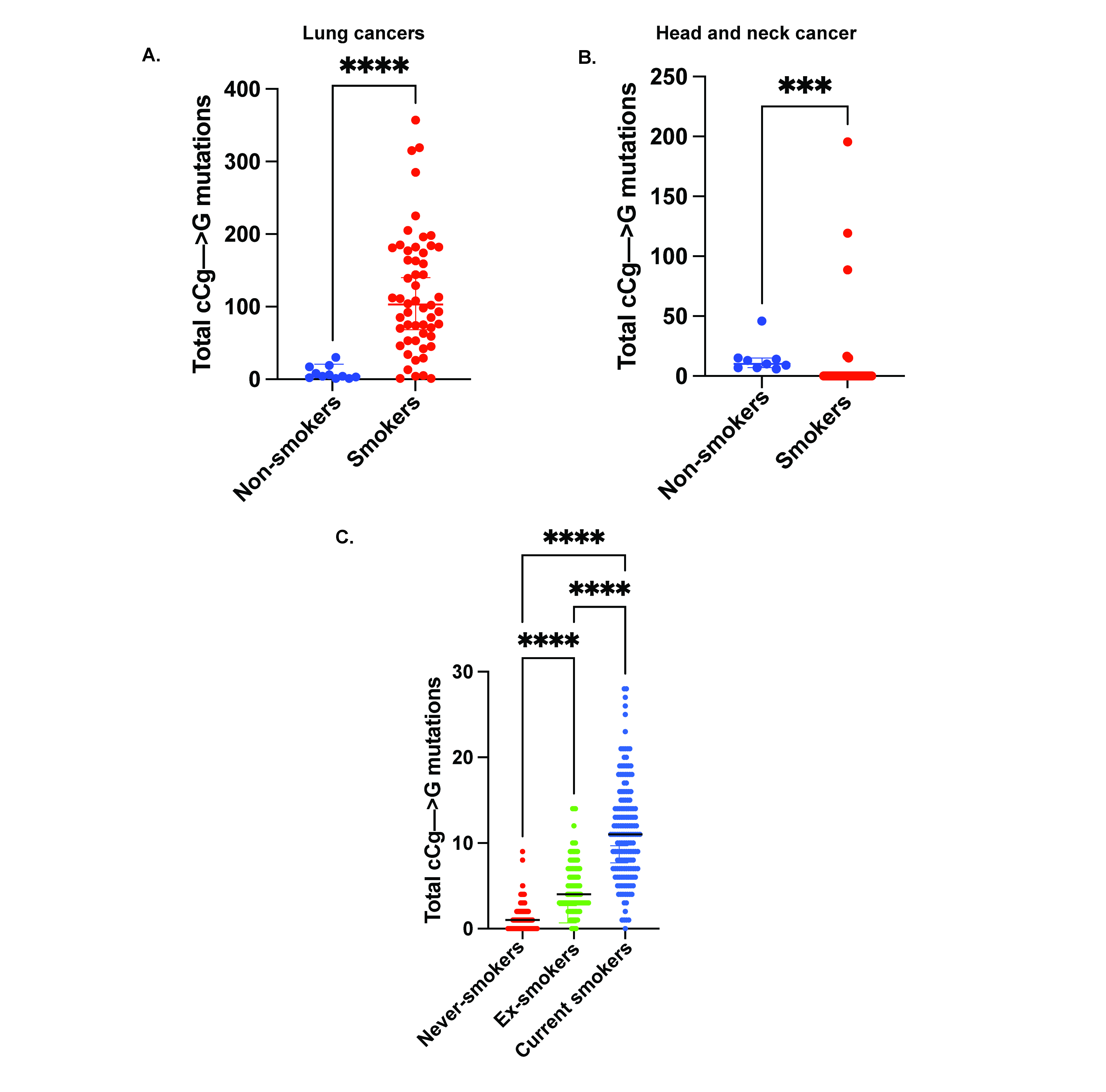
